# Host poly(A) polymerases PAPD5 and PAPD7 provide two layers of protection that ensure the integrity and stability of hepatitis B virus RNA

**DOI:** 10.1101/2021.04.12.439580

**Authors:** Fei Liu, Amy C.H. Lee, Fang Guo, Andrew S. Kondratowicz, Holly M. Micolochick Steuer, Angela Miller, Lauren D. Bailey, Xiaohe Wang, Shuai Chen, Steven G. Kultgen, Andrea Cuconati, Andrew G. Cole, Dimitar Gotchev, Bruce D. Dorsey, Rene Rijnbrand, Angela M. Lam, Michael J. Sofia, Min Gao

## Abstract

Noncanonical poly(A) polymerases PAPD5 and PAPD7 (PAPD5/7) stabilize HBV RNA via the interaction with the viral post-transcriptional regulatory element (PRE), representing new antiviral targets to control HBV RNA metabolism, HBsAg production and viral replication. Inhibitors targeting these proteins are being developed as antiviral therapies, therefore it is important to understand how PAPD5/7 coordinate to stabilize HBV RNA. Here, we utilized a potent small-molecule AB-452 as a chemical probe, along with genetic analyses to dissect the individual roles of PAPD5/7 in HBV RNA stability. AB-452 inhibits PAPD5/7 enzymatic activities and reduces HBsAg both *in vitro* (EC_50_ ranged from 1.4 to 6.8 nM) and *in vivo* by 0.93 log10. Our genetic studies demonstrate that the stem-loop alpha sequence within PRE is essential for both maintaining HBV poly(A) tail integrity and determining sensitivity towards the inhibitory effect of AB-452. Although neither single knock-out (KO) of *PAPD5* nor *PAPD7* reduces HBsAg RNA and protein production, *PAPD5* KO does impair poly(A) tail integrity and confers partial resistance to AB-452. In contrast, *PAPD7* KO could not result in any measurable phenotypic changes, but displays a similar antiviral effect as AB-452 treatment when *PAPD5* is depleted simultaneously. *PAPD5/7* double KO confers complete resistance to AB-452 treatment. Our results thus indicate that PAPD5 plays a dominant role in stabilizing viral RNA by protecting the integrity of its poly(A) tail, while PAPD7 serves as a second line of protection. These findings inform PAPD5 targeted therapeutic strategies and open avenues for further investigating PAPD5/7 in HBV replication.

**Importance:** Chronic hepatitis B affects more than 250 million patients and is a major public health concern worldwide. HBsAg plays a central role in maintaining HBV persistence and as such, therapies reducing HBsAg have been extensively investigated. PAPD5/7 targeting inhibitors, with oral bioavailability, represent an opportunity to reduce both HBV RNA and HBsAg. Here we uncover that the SLα sequence is required for HBV poly(A) tail integrity and RNA stability, and that the antiviral activity of AB-452 mimics the SLα mutants. Although PAPD5 and PAPD7 regulate HBV RNA stability, it remains unclear how they coordinate in stabilizing HBV RNA. Based on our studies, PAPD5 plays a dominant role to stabilize viral RNA by protecting the integrity of its poly(A) tail, while PAPD7 serves as a backup protection mechanism. Our studies may point out a direction towards developing PAPD5-selective inhibitors that could be used effectively to treat chronic hepatitis B.

## Introduction

Globally, more than 250 million patients are chronically infected with hepatitis B virus (HBV) (World Health Organization), but a functional cure of chronic hepatitis B (CHB) is rarely achieved even after years of treatment with nucleos(t)ide analogues (NAs) such as entecavir (ETV) and tenofovir disoproxil fumarate (TDF)(1). Pegylated IFN-α enhances antiviral immune response, but the cure rate remains low and side effects are often difficult to tolerate (2, 3). The major obstacles to curing CHB include the persistence of the episomal covalently closed circular DNA (cccDNA), and an immune system that is tolerized to HBV, likely due to the excess amount of circulating Hepatitis B surface antigen (HBsAg) levels (4–6).

The HBV envelope proteins preS1, preS2 and HBsAg are synthesized in the endoplasmic reticulum and are secreted as both viral and subviral particles (7, 8). HBV virions are double-shelled particles with an outer lipoprotein bilayer containing the envelope proteins, and an inner nucleocapsid that encloses the HBV DNA and viral polymerase. The subviral particles devoid of nucleocapsids and HBV DNA (9, 10) are up to 100,000-fold in excess relative to the virions in the blood of infected patients (11). Such high levels of subviral particles are believed to play a key role in immune tolerance and maintenance of persistent HBV infection (5, 6). In patients with chronic hepatitis B, HBV-specific T cells are depleted or functionally impaired (12–15), and circulating and intrahepatic antiviral B cells are defective in the production of antibodies against HBsAg, with an expansion of atypical memory B cells (16, 17). HBsAg has also been linked to the inhibition of innate immunity and functionality of other immune cell types (18). Therefore, antiviral strategies that aim to target the HBV RNA transcripts could suppress HBsAg production and may break the immune tolerance state to potentially increase the functional cure rate.

Regulation of HBV RNA metabolism involves the post-transcriptional regulatory element (PRE), which is a stretch of ribonucleotides spanning positions 1151-1582 on the viral transcripts that is essential to HBV subgenomic RNA (sRNA) nuclear export and regulation of pregenomic RNA (pgRNA) splicing (19–22). The PRE contains three sub-elements, PREα, PREβ1 and PREβ2. Each sub-element is sufficient to support sRNA nuclear export and HBsAg production, but all three together exhibit much greater activity (23, 24). The PRE is complexed with several RNA binding proteins, including T-cell intracellular antigen 1, La protein, polypyrimidine tract binding protein, ZC3H18 and ZCCHC14 (25–31). These PRE binding proteins may serve to regulate the export and stability of HBV RNAs. In particular, the CAGGC pentaloop sequence/structure of stem-loop alpha (SLα) within the PREα sub-element has been predicted to bind sterile-alpha-motif domain containing proteins (24). Recently, ZCCHC14 (a sterile-alpha-motif containing protein), together with PAPD5 and PAPD7 (the non-canonical poly(A) RNA polymerase associated domain containing proteins 5 and 7), were identified as the cellular binding proteins that interacted with the HBV SLα sequence (32).

The small-molecule compound, RG7834, targets PAPD5/7 and destabilizes HBV RNAs (33–36). Using a genome-wide CRISPR screen, it was subsequently observed that *ZCCHC14* and *PAPD5* were essential for the antiviral activity of RG7834 (31). Interestingly, individual knockdown of *PAPD5* or *PAPD7* had minimal effect against HBsAg production, while knockdown of *ZCCHC14* or double knockdown of *PAPD5*/*7* had a profound anti-HBsAg activity similar to that observed when cells were treated with RG7843 (31, 36). It was further demonstrated that double knockout of *PAPD5/7* reduced guanosine incorporation frequency within HBV RNA poly(A) tails, leading to a proposed model in which HBV RNA recruits the PAPD5/7-ZCCHC14 complex via the CNGGN pentaloop of PRE SLɑ to enable the extension of mixed tailing on HBV poly(A) tails, which subsequently protects the viral RNAs from cellular poly(A) ribonucleases (32).

To gain further insights into how small-molecule inhibitors destabilize HBV RNAs, mechanistic studies were performed using AB-452, an analogue of RG7834, to evaluate its effect in HBV replicating cells and in cells transfected with constructs containing mutations within the PRE sequence. To better understand how PAPD5 and PAPD7 coordinate in the protection of HBV RNAs, both HBV RNA transcripts and their poly(A) tails were analyzed in cells with *PAPD5* and/or *PAPD7* knockout. Our results reveal that HBV utilizes two layers of protection mechanism provided by PAPD5 and PAPD7 to protect their poly(A) tail integrity and RNA stability.

## Results

### AB-452 inhibits HBV *in vitro* and *in vivo*

AB-452 and RG7834 both belong to the dihydroquinolizinones chemical class. The antiviral activities of AB-452 and its diastereomer ARB-169451 were evaluated using multiple *in vitro* HBV replication models including HepG2.2.15 cells (constitutively express HBV through the integrated viral genome), PLC/PRF/5 cells (a patient-derived hepatocellular carcinoma cell line only expressing HBsAg), and HBV infected HepG2-NTCP cells or primary human hepatocytes (PHH) in which viral replication was dependent on cccDNA transcription (Table 1). AB-452 reduced HBsAg, HBeAg, and HBV DNA production with EC_50_ values ranging from 0.28 to 6.8 nM, while its diastereomer ARB-169451 was more than 1,000-fold weaker towards HBsAg inhibition when compared to AB-452 (Table 1). AB-452 antiviral activity was specific for HBV as the compound was inactive against a panel of ten different RNA and DNA viruses with EC_50_ values of >50 μM (Table S1). In addition, the cytotoxicity of AB-452 was evaluated in several cell lines from different tissue origins showing CC_50_ values of >30 μM (the highest concentration tested) (Table S2), demonstrating the selectivity of AB-452.

**Table 1.**
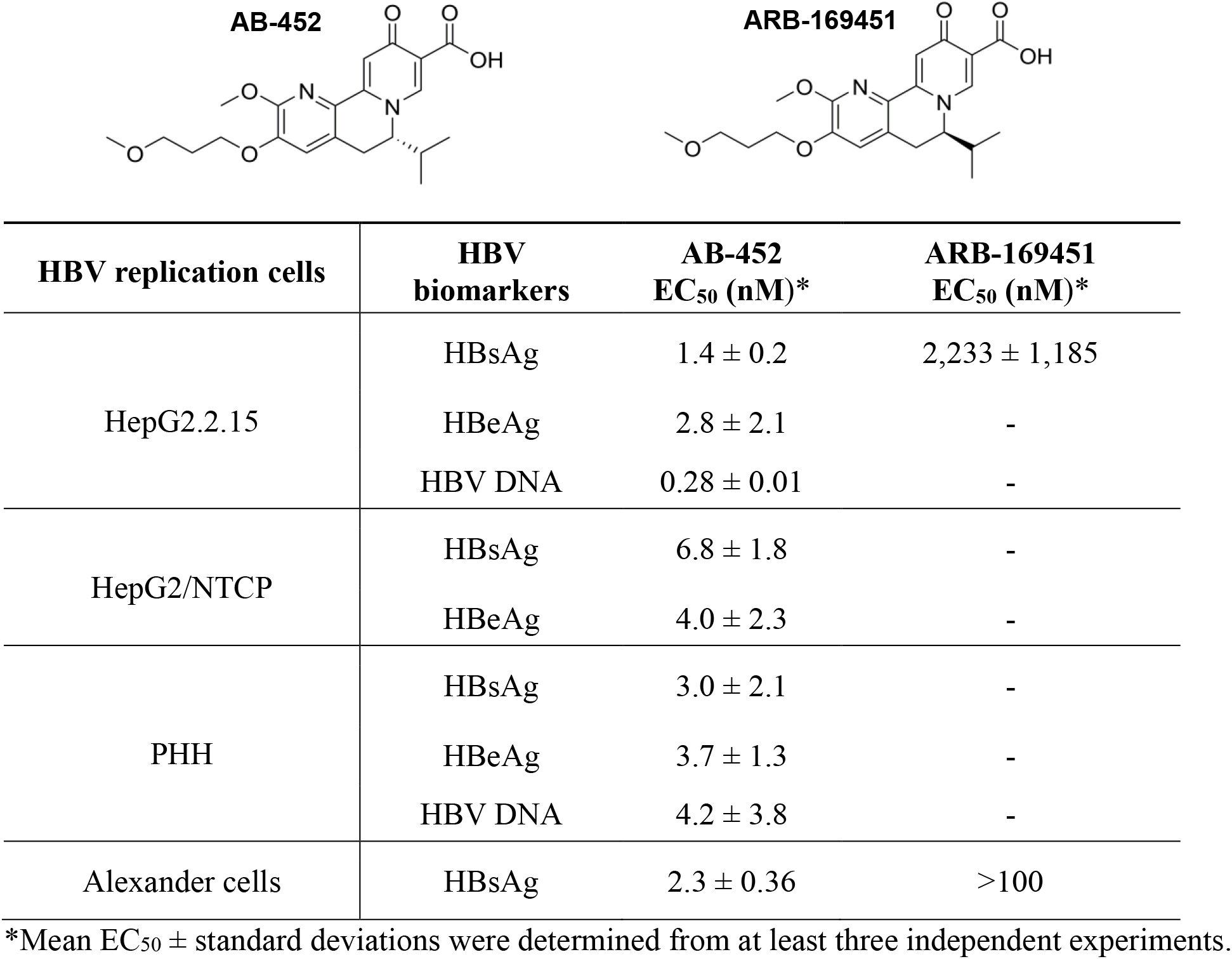
*In vitro* anti-HBV effect of AB-452 and ARB-169451.

To evaluate the effects of AB-452 against the different stages of the viral life cycle, HBV replication intermediates and viral proteins were analyzed from HepG2.2.15 cells treated with AB-452 at a concentration of 50-fold above its EC_50_ value (Fig. 1). The nucleoside analog ETV and two classes of HBV capsid inhibitors GLS-4 (class I) and compound A (class II) (cmpdA chemical structure, Fig. S2) were included as controls targeting the polymerase and core/capsid proteins, respectively. ETV strongly inhibited HBV DNA replication, but it did not reduce other replication intermediates. Consistent with their mechanism of action, capsid inhibitors inhibited pgRNA encapsidation and HBV DNA replication, but had no effect against total pgRNA and sRNA transcripts. On the contrary, AB-452 displayed a unique antiviral phenotype reducing intracellular pgRNA, sRNA, core protein, native capsids, encapsidated pgRNA, and replicating HBV DNA (Fig. 1). Furthermore, the effect of AB-452 against intracellular pgRNA and sRNA was dose dependent and reached a plateau starting at about 100 nM AB-452, with approximately 25% pgRNA and 18% sRNA remaining detectable at the highest concentration tested (1 μM) (Fig. S3A). Results from the time course studies showed that AB-452 induced reduction of pgRNA and sRNA starting at 8 h post treatment and the levels continued to decline through the 48 h treatment period (Fig. S3B).

**Fig. 1.**
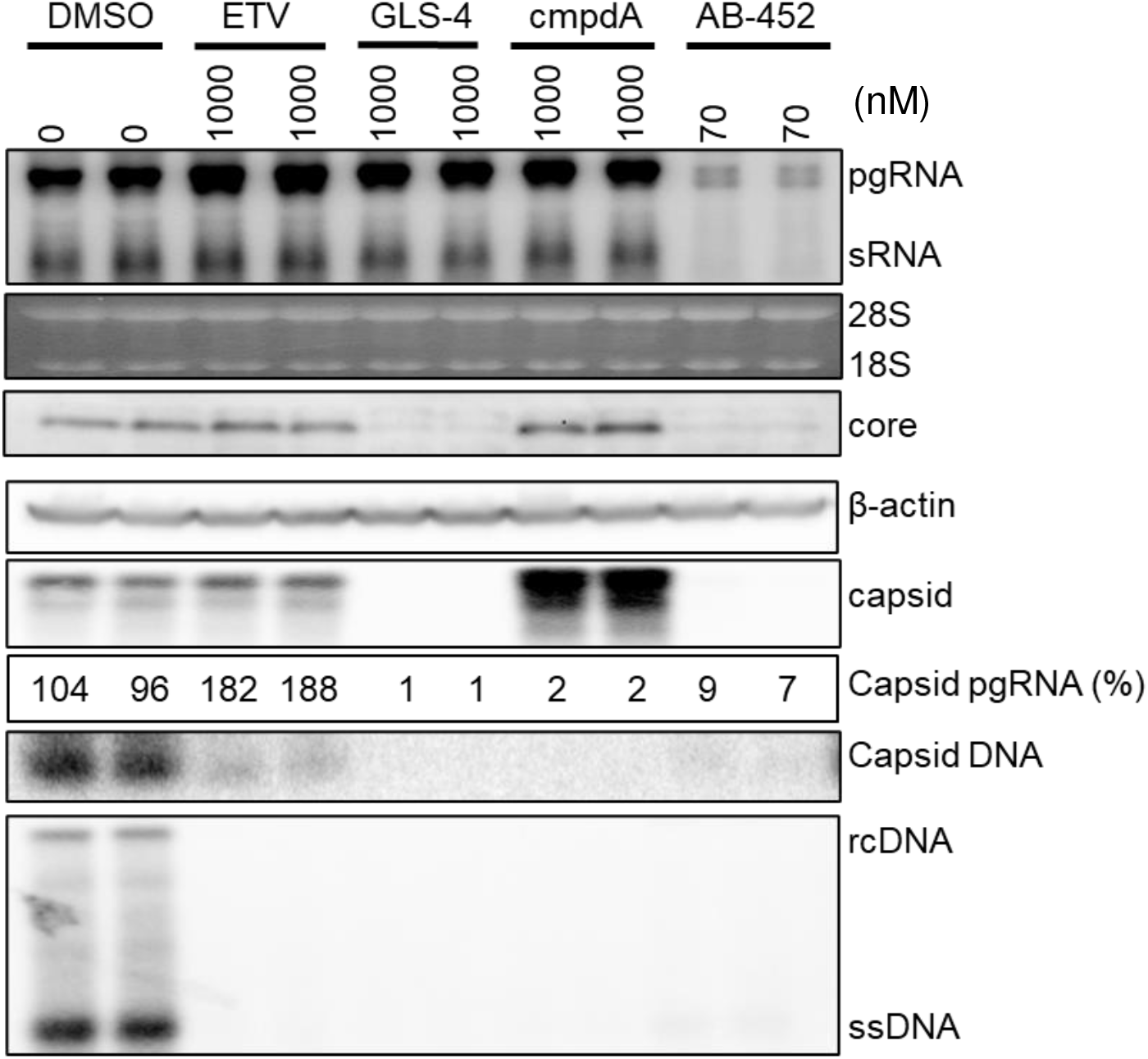
AB-452 interferes with multiple steps of HBV life cycle. HepG2.2.15 cells were treated with DMSO, ETV (1 μM), GLS-4 (1 μM), cmpdA (1 μM), or AB-452 (70 nM) for 6 days. HBV replication intermediates and host markers were analyzed by gel-based analysis. Encapsidated pgRNA (capsid pgRNA) was quantitated by qRT-PCR and expressed as percentage of untreated controls (DMSO).

An AAV-HBV-transduced mouse model was used to assess the anti-HBV effect of AB-452 *in vivo* (Fig. S1). Compared to the vehicle control, oral administration of AB-452 for 7 days at 0.1, 0.3, and 1 mg/kg twice daily resulted in mean 0.68, 0.72 and 0.93 log10 reduction of serum HBsAg (Fig. S1A) and mean 0.79, 1.16 and 0.94 log10 reduction of serum HBV DNA (Fig. S1B), respectively. Inhibition of circulating HBV markers at 0.1, 0.3 and 1 mg/kg doses was found to be correlated with dose-dependent reductions of viral products in the liver: intrahepatic HBsAg levels were reduced by 64, 69 and 83% (Fig. S1C), intrahepatic total HBV RNA levels were reduced by 35, 55, and 66%, and intrahepatic pgRNA levels were reduced by 43, 55, and 63% (Figs. S1D and S1E), respectively. AB-452 treatments were well-tolerated, with no significant change or reduction in body weight in mice throughout the course of the compound treatment compared to those receiving the vehicle control (Fig. S1F). The *in vitro* observation that AB-452 suppressed intracellular HBV RNA was therefore translatable to the *in vivo* AAV-HBV transduced mouse model when treated with AB-452.

### AB-452 promotes HBV RNA degradation through inhibiting PAPD5/7 and blocking guanosine incorporation within HBV poly(A) tails

To investigate the molecular mechanism of how AB-452 inhibits HBV RNA, studies were performed using HepAD38 cells in which HBV transcription is under tetracycline (Tet) regulation. Tet was first removed to induce transcription and accumulation of viral RNAs, and the capsid inhibitor GLS-4 was added to prevent pgRNA encapsidation. Six days later, Tet was added back to shut down further transcription and cells were treated with both GLS-4 and AB-452 for an additional 16 h. The effect of AB-452 on HBV transcripts was evaluated by collecting cells at 0, 2, 4, 8 and 16 h post treatment, and decay of the transcribed HBV RNA was monitored by Northern blot analysis. In the absence of AB-452, HBV RNA levels reduced over time due to natural decay (Fig. 2A). In the presence of AB-452, both pgRNA and sRNA exhibited faster migration starting at 2 h post-treatment and their levels were significantly reduced at 8 and 16 h post-treatment (Fig. 2A). Determination of the pgRNA half-lives (T_½_) showed that AB-452 treatment reduced the T_½_ values from 4.5 h to 2.4 h compared to those from untreated cells (Fig. 2B).

**Fig. 2.**
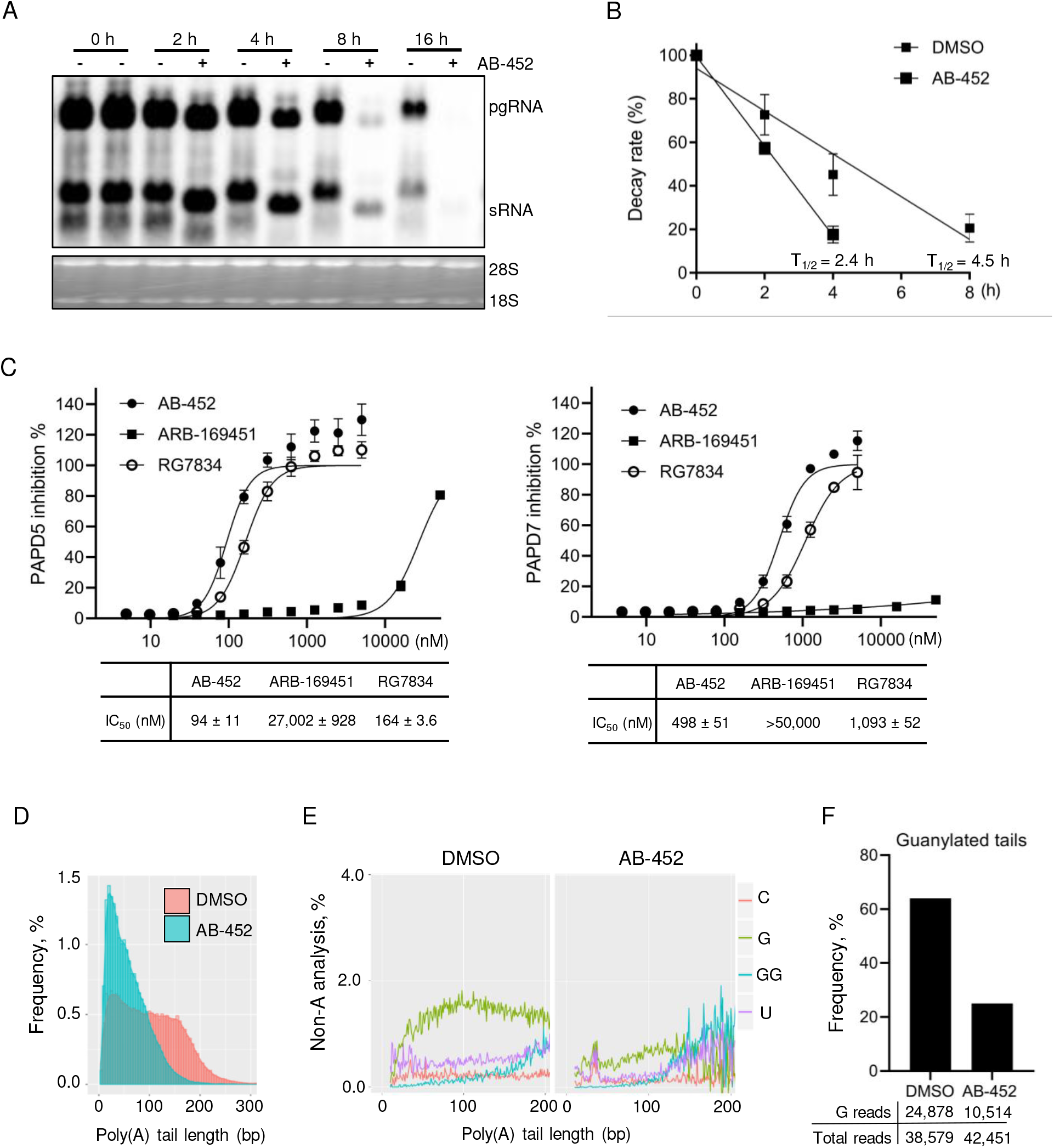
AB-452 promotes HBV RNA degradation through inhibiting PAPD5 and PAPD7 enzymatic activities and blockage of guanosine incorporation into viral RNA poly(A) tails. HepAD38 cells were cultured in the absence of Tet to promote HBV transcription, and the capsid inhibitor GLS4 was included to prevent pgRNA encapsidation for 6 days. On day 7, Tet was added back with media containing either DMSO or AB-452 (70 nM), and cells were harvested either before treatment (time 0 h) or at 2, 4, 8, and 16 h post-treatment. (A) HBV mRNA was analyzed by Northern blot, with ribosomal RNAs as loading control. (B) Decay rate of HBV pgRNA in the presence or absence of AB-452, with calculated T_1/2_ labeled for each treatment (*n* = 2). (C) Effect of AB-452, RG7834, and AB-169451 on the enzymatic activity of PAPD5 and PAPD7. Half-maximal inhibition (IC_50_) for the three compounds were determined based on the dose response curves and reported in the table below each figure. Mean values (± standard derivations) are presented from duplicate experiments. (D) HepAD38 cells were treated with AB-452 (70 nM) for 4 h prior to isolation of intracellular RNA. HBV RNA poly(A) tails were converted into cDNA and amplified for sequencing and tail lengths analysis. (E) Frequency of non-A modifications (C, cytidylation, G/GG, guanylation; U, uridylation) was analyzed within the HBV poly(A) tails from cells treated with or without AB-452 (70 nM). Guanosines were often clustered; tandem GG analysis was made to reflect this observation. (F) Determination of G nucleotide frequency located within the HBV poly(A) tails from cells treated with or without AB-452 (70 nM). The numbers of guanylated tail reads and total tail reads obtained from the NSG sequencing were indicated under each sample. The frequency of guanylated tails of viral mRNAs was calculated for poly(A) tail length of ≥ 10 nt.

Destabilization of HBV RNA by RG7834 was reported to be mediated through inhibiting the PAPD5 and PAPD7 proteins (31, 36). We determined the effect of AB-452 on the enzymatic activity of recombinant PAPD5 and PAPD7 using an ATP depletion biochemical assay (Fig. 2C). Results showed that AB-452 efficiently inhibited PAPD5 with an IC_50_ of 94 nM (Fig. 2C). RG7834 also inhibited PAPD5, although the potency (IC_50_ = 167 nM) was lower than previously reported (IC_50_ = 1.3 nM) (31), possibly due to the more truncated PAPD5 form that was used in the current study. AB-452 and RG7834 inhibited PAPD7 enzymatic activity with IC_50_ values of 498 nM and 1093 nM, respectively. In contrast, the enantiomer ARB-169451 was unable to effectively inhibit PAPD5 (IC_50_ = 27,000 nM) or PAPD7 (IC_50_ >50,000 nM) (Fig. 2C).

RNA metabolism in most eukaryotic mRNAs employs the 3’ deadenylation pathway, in which poly(A) tail shortening is often observed prior to mRNA degradation (37–39). We therefore determined the HBV poly(A) tail length and composition from HepAD38 cells in the presence or absence of AB-452. To amplify the HBV poly(A) tail, G/I (guanosine and inosine nucleotides) tailing was added to the 3’-ends of mRNA transcripts and the newly added G/I tails were used as the priming sites to synthesize the cDNA that would be used for amplification of HBV poly(A) tails. The lengths and compositions of the amplicons containing the HBV RNA poly(A) tails were analyzed by next generation sequencing (PacBio Sequel Sequencing platform). Results showed that majority of the HBV poly(A) tails from untreated samples ranged between 50 to 200 nucleotides in length, with an average tail length of around 100 nucleotides. In contrast, AB-452 treatment reduced the HBV RNA poly(A) tail length by almost 50%, to an average of 58 nucleotides (Fig. 2D and Table S3). The poly(A) tails amplified from β-actin cDNAs served as the negative control, which was not responsive to the treatment (Fig. S4).

It was recently reported that PAPD5/7 extended HBV mRNA poly(A) tails with intermittent guanosine (G), and the incorporation of G could shield them from rapid de-adenylation by cellular deadenylases (32). Since AB-452 inhibited PAPD5/7 enzymatic activities and shortened poly(A) tail lengths, we therefore hypothesized that the G content within the HBV poly(A) tails would be affected by AB-452 treatment. Quantification of the non-adenosine nucleosides within the HBV poly(A) tails indeed revealed that the frequency of G was significantly reduced in the presence of AB-452 (Fig. 2E). The fraction of poly(A) tails containing internal G was reduced from 64% to 25% in the presence of AB-452 compared to those from untreated HepAD38 cells (Fig. 2F). Taken together, the data indicate that inhibition of PAPD5/7 by AB-452 led to blockage of G incorporation and shortening of the poly(A) tail.

### SLα within the PRE sequence is required for HBV RNA integrity and AB-452 susceptibility

We and others have determined that reduction of HBV RNA by RG7834 is dependent on the HBV PRE (31, 32, 35), which partially overlapped with the HBx coding region. To further define the involvement of the sub-elements within PRE on HBV RNA stability and AB-452 susceptibility, several reporter plasmids were constructed (Fig. 3A): 1) H133 is the wild type construct supporting the expression of 2.1 kb HBV sRNA expression, 2) H133_Gluc is derived from H133 but with the HBsAg coding sequence replaced with *Gaussia* luciferase, 3) Gluc_dHBx is derived from H133_Gluc but with most of the HBx coding sequence deleted (nucleotide 1389 to 1991) and the HBV poly(A) replaced with the SV40 poly(A) signal, and 4) Gluc_rcSLα is derived from Gluc_dHBx with an inverted SLα sequence.

**Fig. 3.**
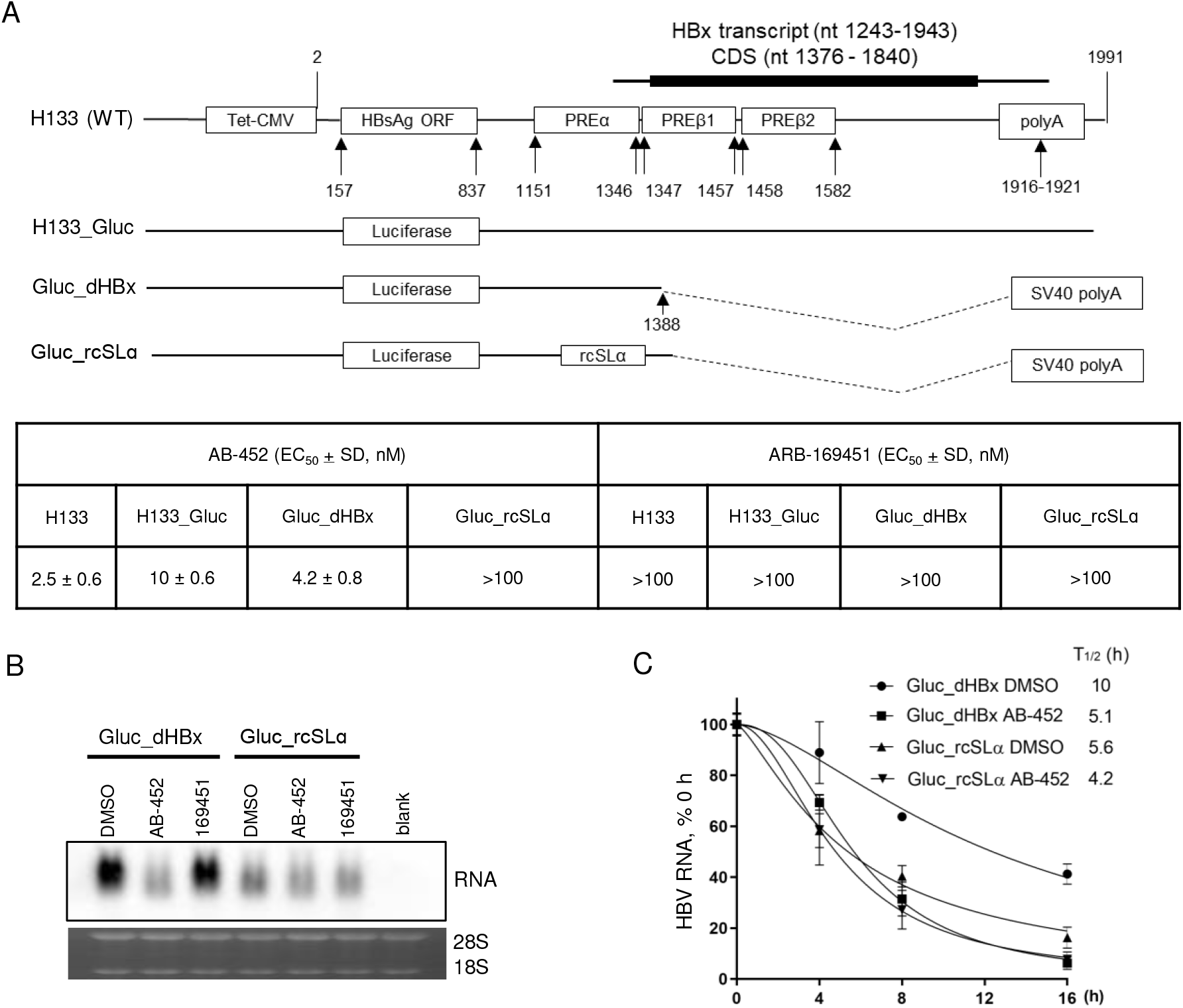
SLα sequence within the HBV PREα sub-element is essential for RNA stability and AB-452 activity. Evaluation of AB-452 against H133 and Gaussia luciferase (Gluc) encoded plasmids containing either wildtype HBV PRE or inversion-derived mutants. (A) Schematic representation of H133 and the Gluc-encoded constructs. Huh-7 cells were transfected with each of these plasmids and susceptibility to AB-452 was evaluated by monitoring HBsAg or Gluc activity. The inactive enantiomer ARB-169451 was included as a negative control. Mean values (± standard derivations) are presented from triplicate experiments. (B) The HBx deletion variants containing either the WT SLα (Gluc_dHBx) or the inverted SLα (Gluc_rcSLα) sequence were transfected into Huh-7 cells, which were treated with DMSO, AB-452 (100 nM) or ARB-169451 (100 nM) for 5 days. Effect of AB-452 against HBV RNA was analyzed by Northern blot with ribosomal RNAs as loading control. (C) Kinetics of HBV RNA degradation in Huh-7 cells transfected with Gluc_dHBx or Gluc_rcSLα plasmids. Transcription proceeded for 2 days prior to the addition of tetracycline with or without AB-452 (100 nM). Cells were harvested before treatment (time 0 h) and at 4, 8, and 16 h post treatment. HBV RNA decays were analyzed by qRT-PCR. Data and error bars represent mean % HBV RNA and standard deviations relative to time 0 of each condition from at least three independent experiments.

AB-452, but not its enantiomer ARB-1694151, inhibited both HBsAg and Gluc expression in cells transfected with H133 (EC50 = 2.5 nM), H133_Gluc (EC50 = 10.0 nM), or the Gluc_dHBx construct (EC50 = 4.2 nM) (Fig. 3A). These data indicate that AB-452 antiviral activity was not dependent on the HBsAg sequence, HBx sequence or the HBV poly(A) signal sequence. On the other hand, inverting the SLα sequence (Gluc_rcSLα) completely abolished sensitivity to AB-452 (EC50 >100 nM) (Fig. 3A and 3B). Interestingly, we observed that the transcribed RNA from the Gluc_rcSLα transfected cells showed reduction in RNA levels and appeared smaller in size when compared to the RNA from cells transfected with the Gluc_dHBx plasmid, with or without AB-452 treatment (Fig. 3B). The rates of HBV RNA decay revealed that AB-452 treatment reduced Gluc_dHBx RNA half-lives (T½) from 10 h to 5.1 h when compared to DMSO treated cells (Fig. 3C). In contrast, the Gluc_rcSLα RNA was unstable (T½ = 5.6 h) and its T½ was only slightly reduced by AB-452 (T½ = 4.2 h) (Fig. 3C).

In addition to SLα, HBV PREα contains another cis-acting element known as La protein binding element, these two cis-acting elements were included in a 109 nucleotides sequence that was critical for RG7834 sensitivity (35). The requirement of these two elements was studied by generating two additional H133 derived constructs, H133_dSLα and H133_dLa, in which the SLα sequence and the La element was deleted, respectively. AB-452 inhibited HBsAg production in H133_dLa transfected cells with similar efficiencies as the wildtype H133 construct (EC_50_ = 4.3 and 2.5 nM, respectively), indicating that the La protein binding element was not essential for susceptibility to AB-452. Alike to the results observed in the Gluc_rcSLα transfection, AB-452 was inactive against the H133_dSLα (EC_50_ >100 nM) (Fig. 4A, Fig. S5). Deleting the SLα sequence also led to the shortening and reduction of sRNA level (Fig. 4B), as well as reduced transcripts T_½_ (Fig. 4C). Cells transfected with H133_dSLα showed reduced sensitivity to AB-452, with transcript half-life being only slightly reduced from 6.2 h to 5.5 h when compared to those treated with DMSO (Fig. 4C).

**Fig. 4.**
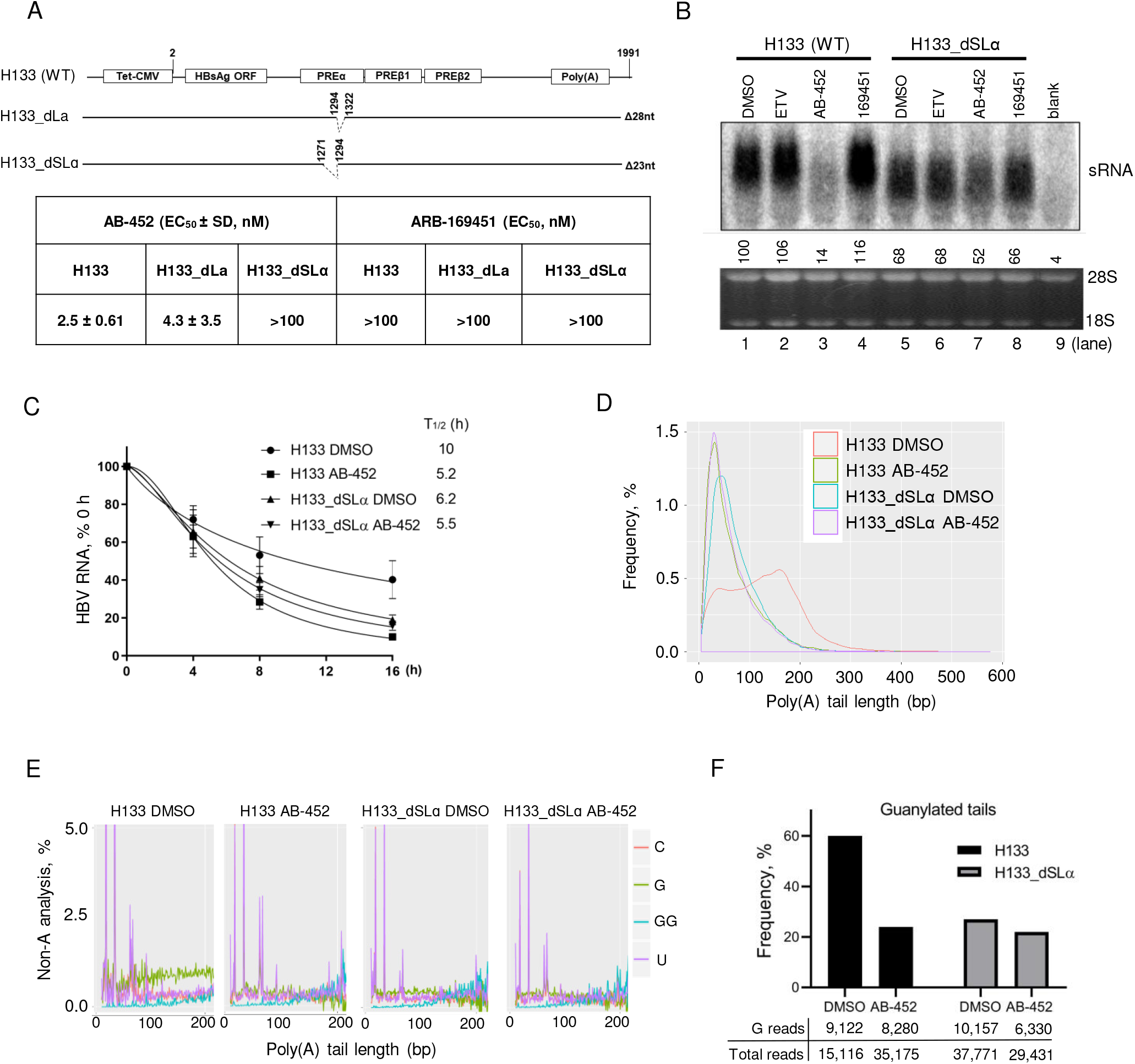
SLɑ determines HBV RNA poly(A) tail integrity and stability. Evaluation of the SLα sequence on sensitivity towards AB-452 and HBV RNA stability. (A) H133_dSLα and H133_dLα were mutants with either SLα (nt 1294-1322) or La binding site (nt 1271-1294) deleted. Huh-7 cells were transfected with H133, H133_dLa or H133_dSLα plasmids and treated with AB-452 for 5 days. The activities of AB-452 and ARB-169451 against HBsAg production were determined and its EC_50_ values summarized in the table below the schematic representation. Mean values (± standard derivations) are determined from triplicate experiments. (B) Levels of HBV sRNA from transfected cells treated with ETV (1 μM), AB-452 (100 nM), and 169451 (100 nM) were analyzed by Northern blot. (C) HBV sRNA was quantitated by qRT-PCR assay, with calculated decay T_1/2_ labeled under each treatment (*n* = 3). (D) HBV RNA poly(A) tails were sequenced and analyzed for frequency of tail lengths from cells transfected with the H133 or H133_dSLα plasmids treated with or without AB-452 (100 nM). (E) Frequency of non-A modifications (G, guanylation; U, uridylation; C, cytidylation) within the ploy(A) tail of HBV mRNAs were analyzed. Tandem GG analysis was performed to analyze clustered guanosines. (F) The frequency of guanylated tails of viral mRNAs was calculated with a poly(A) tail length of ≥ 10 nt. The numbers of guanylated tail reads and total tail reads obtained from the NSG sequencing were indicated under each sample.

NGS analysis of the sRNA poly(A) tails showed that AB-452 reduced the average poly(A) tail length of H133 transcripts from 124 to 64 nucleotides (Fig. 4D). The poly(A) tails from the H133_dSLα transcripts were 62 and 71 nucleotides with and without AB-452 treatment, respectively (Fig. 4D). In terms of the poly(A) tail composition, the guanylation frequency was highest in cells transfected with the wildtype PRE (H133), and the overall guanylation frequency was reduced from about 60% to 24% in the presence of AB-452 (Fig. 4E and 4F). In contrast, the guanylation frequency in the H133_dSLα transcripts already appeared low (22% to 27%) with and without AB-452 treatment (Figs. 4E and 4F). Taken together, these data provide first line evidence demonstrating that the SLα sequence serves to stabilize the viral transcripts through maintaining the poly(A) tail lengths and mixed-nucleotides composition.

### PAPD5 and PAPD7 determine HBV RNA integrity and stability

To understand the individual role of PAPD5 and PAPD7 in regulating HBV RNA stability and poly(A) tail integrity, knockout (KO) cell lines with deletion of *PAPD5* (*P5*_KO), *PAPD7* (*P7*_KO), or both *PAPD5/7* (double knockout, *P5/7*_DKO) were isolated using CRISPR-Cas9 gene editing and HepG2-NTCP cells. It was reported that ZCCHC14 (Z14), which interacts with PAPD5/7 and the HBV PRE, plays an important role in maintaining HBV RNA integrity and stability (32). *Z14* KO cell lines (*Z14*_KO) were generated to assess the involvement of Z14 on regulating HBV RNA. In addition to the parental wildtype (WT) HepG2-NTCP cell line, two additional WT cell clones (T3-4 and T2-14) were included as clonal controls. The full-allelic KO genotype for all the individual cell clones was confirmed by DNA sequencing (Fig. S7A), and *PAPD5* and *Z14* knockout were also confirmed at the protein level (Fig. S7B). PAPD7 protein expression could not be evaluated by Western blot due to the lack of an efficient PAPD7 specific antibody, but the PAPD7 KO genotype was confirmed by DNA sequencing (Fig. S7A).

Overall, cell proliferation analysis suggests that PAPD5, PAPD7, and Z14 were not critical for cell survival (Fig. 5A). The effect of knocking out *PAPD5/7* and *Z14* on viral protein production and HBV replication was examined by using two independent systems: adenovirus-encoded HBsAg transduction and HBV infection (Figs. 5B-C). In the adenovirus transduction studies, single KO of *PAPD5* or *PAPD7* did not reduce HBV replication or antigens production compared to the parental cell clones. HBsAg expression in the *P5/7*_DKO and *Z14*_KO clones was about 50% lower than that of the WT or *PAPD5/7* single KO clones in the 5 days culture (Fig. 5B). In the HBV infection studies, the levels of viral proteins and HBV DNA were much lower in the *P5/7*_DKO and *Z14*_KO clones compared to the WT or *PAPD5/7* single KO clones in the 9 days culture (Fig. 5C).

**Fig. 5.**
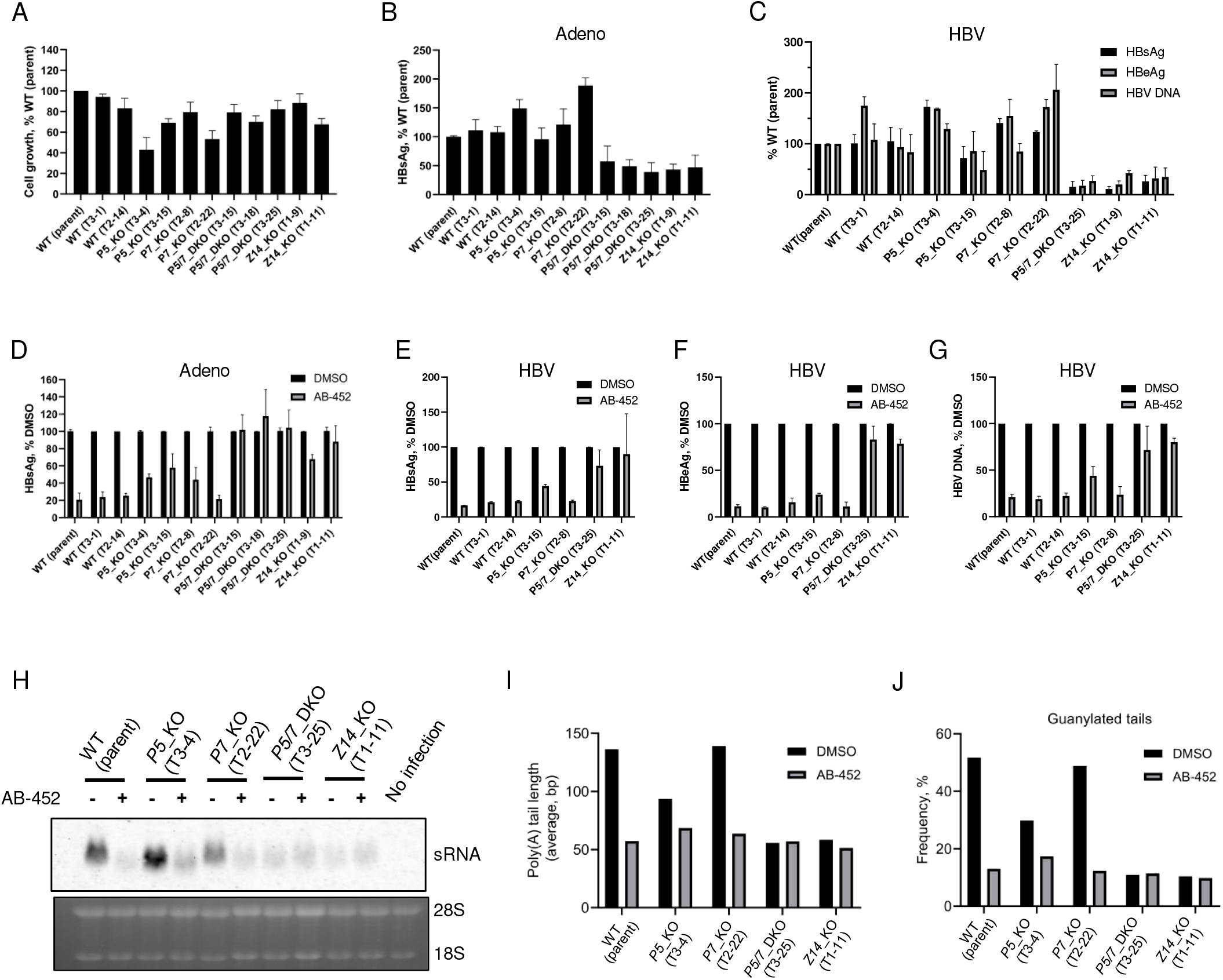
Knockout of *PAPD5/7* and *ZCCHC14* destabilizes and desensitizes HBV RNA to AB-452. *PAPD5*, *PAPD7*, *ZCCHC14*, or both *PAPD5* and *PAPD7* were knocked out in HepG2-NTCP cells by CRISPR-Cas9 and gRNAs designed to target these genes. (A) Cell proliferation analysis of *PAPD5*, *PAPD7* and *Z14* KO or WT clones was analyzed. Percentage cell growth relative to the WT parent HepG2-NTCP cells was determined for each tested clone. (B) Adenoviruses carrying HBsAg coding sequence were used to transduce either WT, *PAPD5, PAPD7* or *ZCCHC14* KO cell clones, extracellular HBsAg was measured on day 5 post transduction. Percentage of HBsAg relative to the WT parent HepG2-NTCP cells was determined. (C) HepG2-NTCP cells were infected with HBV inoculum. HBsAg, HBeAg and HBV DNA were measured on day 9 post infection in the KO clones and normalized to the WT parent cells. (D) AB-452 activity of HBsAg inhibition was evaluated in the *PAPD5/7* single or double KO and *ZCCHC14* KO clones infected with adenoviruses. (E-G) AB-452 antiviral activity was evaluated in HBV infected HepG2-NTCP clones. (H) HBV sRNA was analyzed by Northern Blot in the *PAPD5/7* single or double KO and *ZCCHC14* KO cell clones treated with and without AB-452 for 5 days. (I) HBV sRNA poly(A) tails were sequenced for the analysis of tail lengths and (J) guanylation incorporation frequency. The Mean values and standard derivations were plotted at least from duplicate experiments for the Figs. A-G.

We next examined the impact of deleting *PAPD5*, *PAPD7*, *Z14*, or both *PAPD5/7* on compound sensitivity. In adenovirus transduced cells, AB-452 inhibited HBsAg production from WT and *P7*_KO cells with similar EC_50_ values of 9 nM and 10 nM, respectively. However, AB-452 was about 7-fold less active against the *P5*_KO cells (EC_50_ = 72 nM) (Table 2). Susceptibility to AB-452 was also evaluated using HBV infected HepG2-NTCP cells: results showed that AB-452 inhibited WT and *P7*_KO cells with similar efficiencies (EC_50_ values of 11 nM and 9 nM, respectively), but was again less active against the *P5*_KO cells (EC_50_ = 85 nM) (Table 2). A similar trend was also observed with RG7834, suggesting this differentiated antiviral activity was not AB-452 specific. The antiviral data are consistent with the finding that AB-452 and RG7834 were more efficient against PAPD5 than PAPD7 in the enzymatic assays (Fig. 2C). Among the *P5/7*_DKO and *Z14*_KO cell lines, AB-452 treatment did not show further inhibition compared to the untreated controls (Figs. 5D to 5G).

**Table 2.** Anti-HBV effect of AB-452 and RG7834 in *PAPD5* or *PAPD7* KO cell lines.

| Cell lines | Compound | Adeno infection |  | HBV infection |  |
| --- | --- | --- | --- | --- | --- |
|  |  | EC<sub>50</sub> (nM) | #FC vs WT | EC<sub>50</sub> (nM) | FC vs WT |
| WT (parent) | AB-452 | 9.0 ± 4.6 | 1 | 2.7 ± 1.8 | 1 |
|  | RG7834 | 11.8 ± 9.6 | 1 | - | - |
| *P5_KO* (T3-4) | AB-452 | 71.6 ± 31.2 | 8.0 | 37.7 ± 8.3 | 14.0 |
|  | RG7834 | 126 ± 74.7 | 10.7 | - | - |
| *P5_KO* (T3-15) | AB-452 | 56.0 ± 34.0 | 6.3 | 39.3 ± 3.2 | 14.6 |
|  | RG7834 | 85.7 ± 59.8 | 7.3 | - | - |
| *P7_KO* (T2-8) | AB-452 | 10.0 ± 1.0 | 1.1 | 3.8 ± 1.5 | 1.4 |
|  | RG7834 | 23.7 ± 15.2 | 2.0 | - | - |
| *P7_KO* (T2-22) | AB-452 | 7.1 ± 1.3 | 0.8 | 3.2 ± 1.9 | 1.2 |
|  | RG7834 | 10.8 ± 2.1 | 0.9 | - | - |

Mean EC<sub>50</sub> ± standard deviations were determined from three independent experiments. #FC: fold change.

Interestingly, while there was no appreciative reduction of HBV protein and DNA production observed from the *P5*_KO cells (Figs. 5B-C), we noted that the sRNA migrated faster than those from the WT and *P7*_KO cells (Fig 5H). Intrigued by this observation, the sRNA from WT and the various KO cells were further characterized by NGS analysis. Results revealed that knocking out *PAPD5* alone, but not *PAPD7*, reduced both the poly(A) tail lengths (from >136 bp to 94 bp) and guanylation frequency of sRNA (from ~50% to 30%) when compared to the WT cells (Figs. 5I, 5J and S8, and Table S5). AB-452 treatment led to reduction of poly(A) tail length (from >136 bp to ~60 bp) and guanylation (from ~50% to ~10%) in both WT and *P7*_KO cells. The *P5*_KO cells appeared less sensitive to AB-452 in its shortening of poly(A) tail lengths and guanosine incorporation. Cells with *Z14*_KO and *PAPD5/7*_DKO already showed drastically reduced levels of sRNA, poly(A) tail lengths (51 - 58 bp) and guanosine incorporation (~10%), with and without AB-452 treatment. Taken together, these results suggest that of the two noncanonical poly(A) polymerases, PAPD5 appeared to play a major role in determining viral poly(A) tail integrity, guanosine incorporation and AB-452 sensitivity.

## Discussion

Current therapies for chronic hepatitis B patients rarely achieve functional cure, which is characterized as sustained loss of HBsAg with or without HBsAg antibody seroconversion (40). The discovery of RG7834 has raised significant interest as this class of small-molecule inhibitors has the potential to reduce both HBV RNA and viral proteins, which are distinct from direct-acting antivirals targeting the HBV polymerase and capsid proteins (31, 36). AB-452 is an analog of RG7834 with a similarly broad antiviral effect against multiple HBV replication intermediates. It has been appreciated that integrated HBV DNA is a major source of HBsAg expression in HBeAg negative patients (41). Our data indicate that AB-452 can reduce HBsAg produced from cccDNA in HBV-infected cells as well as from integrated HBV DNA in patient-derived hepatocellular carcinoma cells (Table 1). Furthermore, oral administration of AB-452 substantially reduced HBV DNA, HBsAg, HBeAg, and intrahepatic HBV RNA from AAV-HBV-infected mice. Our studies here provide insights into the mode of action for AB-452 and further characterize the RNA stabilization mechanisms utilized by the virus. Our results demonstrate that the cis-acting SLα viral sequence and the trans-acting host factors PAPD5 and PAPD7 coordinate to protect viral RNA. Interference of such viral-host interactions through small-molecule compounds treatment or genetic mutations led to destabilization of viral transcripts and reduction of HBsAg.

The requirement of PAPD5/7 and ZCCHC14 to form a complex with HBV RNA through the PRE element for stabilizing HBV RNA has been described (31, 32). Since the ZCCHC14/PAPD5/7 complex is recruited onto the SLα sequence, it is conceivable that mutating the SLα sequence may disrupt the binding of the ZCCHC14/PAPD5/7 complex and consequently affect HBV RNA stability. Here, our studies provided the genetic evidence that an intact SLα sequence is indeed critical for maintaining HBV poly(A) tail integrity and stability, as inverting or deleting this sequence both destabilize HBV RNA. Notably, the phenotype of the SLα deletion and inversion mutants resembled the antiviral effect of AB-452: cells treated with AB-452 display the phenotypes of HBV poly(A) tail shortening, reduced guanosine incorporation, and HBV RNA degradation.

Initial studies suggest that PAPD5 and PAPD7 may provide redundant if not identical role(s) in protecting HBV RNA stability (31, 32, 36, 42). However, our results from the *P5*_KO and *P7*_KO cell lines would argue that PAPD5 and PAPD7 may serve two lines of protection in maintaining the stability of HBV RNA. *P5*_KO, but not *P7*_KO, impaired poly(A) tail integrity. Moreover, the phenotypic measurements we monitored so far indicate that the *P7*_KO cells were similar to WT cells, further supporting that PAPD5 expression alone could support viral RNA integrity and stabilization (Fig. 5). These data suggest that PAPD7 did not actively contribute to HBV RNA protection in the presence of PAPD5, but instead served as a second line of protection by moderately extending HBV poly(A) tail when PAPD5 was depleted (Figs. 5I and 5J). Results from the enzymatic assays show that PAPD5 was more robust than PAPD7 in the extension of poly(A) tails (Fig. S9), supporting our argument that PAPD5 would be the major host factor in protecting HBV RNA. Immune precipitation experiments conducted by two independent research groups indicated that both PAPD5 and PAPD7 were bound to HBV mRNA, with PAPD7 at a lower level compared to PAPD5(32, 42). Further studies would be required to clarify the role of PAPD7 in HBV RNA metabolism in WT cells.

Another noteworthy observation from this study is that the two HBV RNA destabilizers, AB-452 and RG7834, displayed different inhibitory efficiencies against PAPD5 and PAPD7. Both compounds were 5- to 7-fold less efficient against the enzymatic activities of PAPD7 compared to PAPD5, which was in turn consistent with the results from cell-based studies in which AB-452 and RG7834 displayed a 5- to 10-fold reduction in activities against HBsAg production in the *P5*_KO cells (in which PAPD7 is present) when compared to those from the WT and *P7*_KO cells (in which PAPD5 is present). These data suggest that it may not be critical to completely inhibit PAPD7 to achieve HBV RNA destabilization. However, not all HBV RNA destabilizers differentiate between PAPD5 and PAPD7, as compounds from another chemical class showed no discrimination between PAPD5 and PAPD7 (manuscript in preparation). These data, together with the genetic studies, support the hypothesis that PAPD5 could be more essential than PAPD7 in stabilizing HBV RNA. Our results further suggest that developing PAPD5-selective inhibitors of HBV replication could be pharmacologically feasible.

Here, we propose a working model of the interplay between HBV transcripts and the cellular ZCCHC14/PAPD5/7 RNA metabolism machineries (Fig. 6). Maintenance of HBV RNA stability is a dynamic process regulated by canonical and non-canonical poly(A) polymerases and deadenylases. PAPD5 could form a complex with ZCCHC14, which directs the non-canonical polymerase onto the viral transcripts through the SLα within the HBV PRE sequence. Assembly of the ZCCHC14/PAPD5 onto SLα within the HBV PRE sequence facilitates the addition of G while extending the poly(A) tail. This guanylation process may stall the cellular poly(A) exonuclease and terminate further deadenylation, thus protecting the RNA from degradation. When PAPD5 is depleted, ZCCHC14/PAPD7 complex may bind to HBV RNA and protects its degradation, however PAPD7 is less effective for poly(A) extension and guanylation incorporation. When HBV is challenged by PRE mutations or HBV RNA destabilizers such as AB-452, viral RNA integrity and stability are disrupted due to disarraying or inhibition of the ZCCHC14/PAPD5/7 complex from interacting with the SLα sequence.

**Fig. 6.**
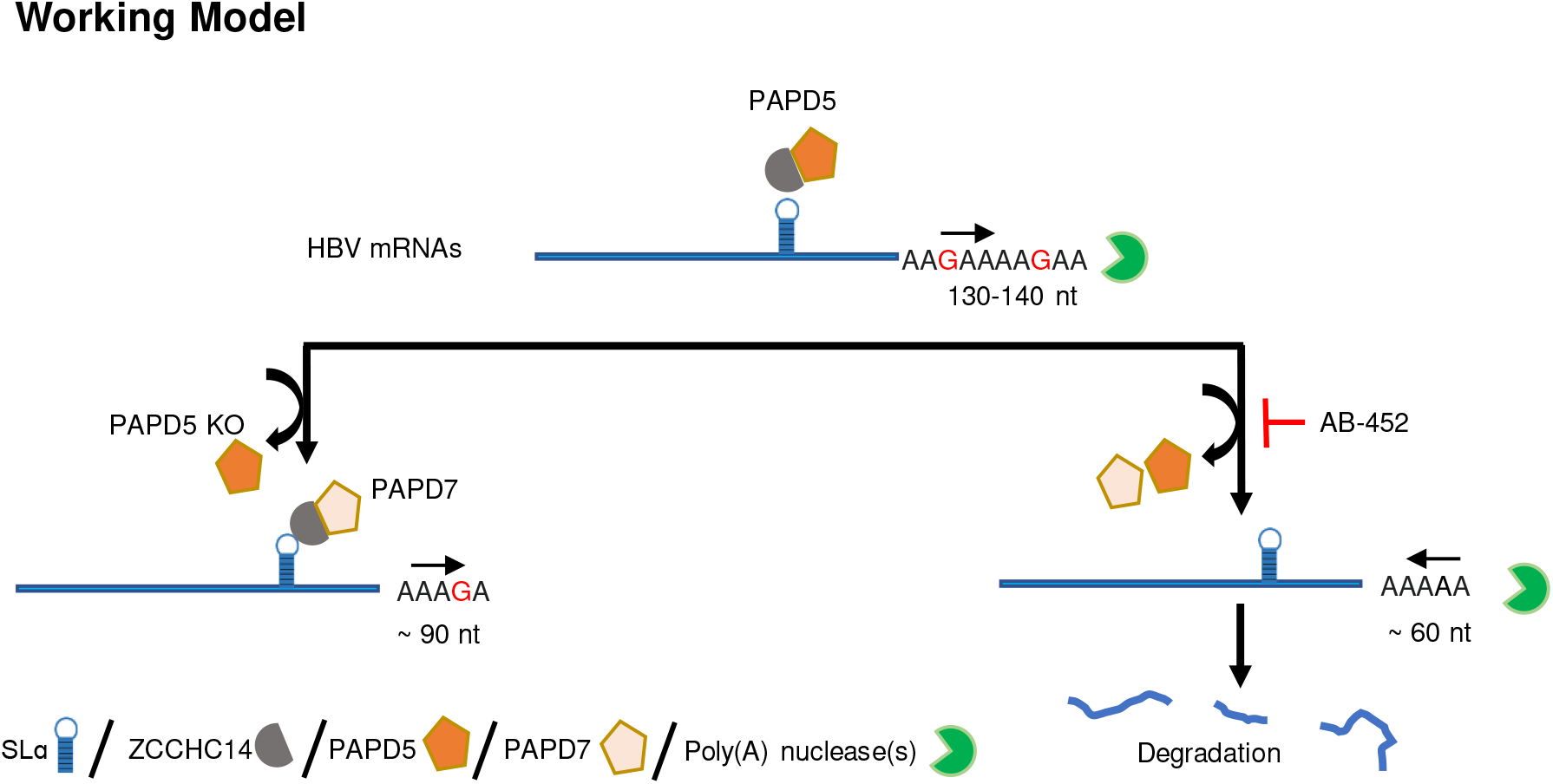
A proposed model illustrating the interplay between HBV RNA cis-elements and the host factors PAPD5 and PAPD7 in maintaining HBV RNA integrity and stability.

## Methods and materials

Detailed methods and materials are provided in the online Supporting information of this paper.

### Cell lines and culture

HepG2.2.15, HepAD38, HepDE19 cells and PLC/PRF/5 cells were cultured in DMEM/F12 medium (Corning, NY, USA), supplemented with 10% fetal bovine serum (Gemini, CA, USA), 100 U/ml penicillin and 100 μg/ml streptomycin. Huh-7 cells (Creative Bioarray, NY, USA) were cultured in RPMI 1640 medium (Basel, Switzerland) containing 10% fetal bovine serum 100 U/ml penicillin and 100 μg/ml streptomycin. HepG2-hNTCP-C4 cells were cultured in DMEM medium (Gibco^TM^, MA, USA) containing 10% fetal bovine serum and 10 mM HEPES (Gibco^TM^). *PAPD5, PAPD7* and *ZCCHC14* were knocked out in the HepG2-hNTCP-C4 cells by using CRISPR technology at GenScript (Piscataway, NJ). Briefly, HepG2-hNTCP-C4 cells were transfected with gRNA and Cas9 expression plasmids. Single cell clones were generated and confirmed by Sanger sequencing (S5A Fig.). The obtained clones were expanded and further evaluated by the Western blots (S5B Fig.). The gRNA sequences are listed as follows: *PAPD5* gRNA 5’- GACATCGACCTAGTGGTGTTTGG-3’, *PAPD7* gRNA 5’-ATATTTGGCA GCT TTAGTACAGG-3’, and *ZCCHC14* gRNA 5’-GCGTGAGACCCGCACCCCCG-3’.

### Measurement of extracellular HBsAg, hepatitis B e antigen (HBeAg) and HBV DNA

HBsAg and HBeAg from the supernatants of cultured HepG2.2.15 cells (43) were measured using the chemiluminescence-based immunoassay per manufacturer’s instructions (AutoBio Diagnostics Co, China). Secreted HBV DNA was extracted according to manufacturer provided protocol (Realtime Ready Cell Lysis Kit, Roche, Mannheim, Germany) and quantified in a qPCR assay (LightCycler® 480 SYBR Green I Master, Roche) with the 5’-GGCTTTCGGAAAATTCC TATG-3’ (sense) and 5’-AGCCCTACGAACCACTGAAC-3’ (antisense) primers using the PCR conditions of denaturing at 95 °C for 5 min, followed by 40 cycles of amplification at 95 °C for 15 s and 60 °C for 30 s.

### PRE cis-elements analysis

Constructs containing either HBsAg or the Gaussia luciferase reporter genes were synthesized (GenScript). The H133 encodes the full HBsAg transcript sequence (spanning nt 2 −1991, U95551) under the regulation of tetracycline controlled CMV promoter. The H133_dSLα and H133_dLa are derived from pH133 with either the SLα sequence (nt 1294 - 1322) or the La protein binding site (nt 1271 - 1294) deleted, respectively. For the luciferase-based plasmids, the HBsAg CDS was replaced with the Gaussia luciferase (Gluc) reporter gene to generate the construct H133_Gluc. To make Gluc_dHBx, the HBx coding sequence was deleted, while the Gluc_rcSLα is derived from Gluc_dHBx with an inverted SLα sequence.

Huh-7 cells were transfected with the HBsAg or luciferase reporter derived plasmids per manufacturer’s instructions (Lipofectamine 3000, Invitrogen, MA, USA). Cells were treated with the indicated compounds for 5 days. Culture supernatants were used for HBsAg or luciferase measurement (Pierce Gaussia Luciferase Glow Assay Kit, ThermoFisher Scientific, Waltham, MA, USA). Cells were collected for HBV RNA transcript and cellular ribosomal RNA analysis by Northern blots.

### Infection of HepG2-hNTCP-C4 cells and primary human hepatocytes (PHH)

HepG2-hNTCP-C4 cells and PHH (BioIVT, Westbury, NY, USA) were cultured in complete DMEM medium containing 2% DMSO overnight, infected with HBV at a MOI of 100-250 GE/cell, and subsequently treated with compounds for 11-16 days with medium and compound treatments refreshed every 2-3 days. The supernatants were harvested for HBsAg and HBeAg analysis (ELISAs, International Immuno-Diagnostics, CA, USA). HBV DNA was extracted from cell lysates per manufacturer’s instructions (Qiagen DNeasy 96 Blood and Tissue Kit, Qiagen, Hilden, Germany). HBV DNA was detected by qPCR using primers and probe as follows: 5’- GTCCTCAAYTTGTCCTGG-3’ (sense), TGAGGCATAGCAGCAGGA-3’ (antisense), and Probe /56-FAM/CTGGATGTGTCT GCGGCGTTTTATCAT/36-TAMSp/.

### Northern and Southern blots

Northern and Southern blots were performed as described previously (44). Total intracellular RNA samples were separated in 1.5% agarose gels, transferred onto Hybond-XL membrane and probed with α-^32^P-UTP (Perkin Elmer, CT, USA) labeled HBV plus-strand-specific riboprobe. Intracellular viral DNA was analyzed by Southern blot hybridization with an α-^32^P-UTP labeled HBV minus-strand-specific riboprobe. Membranes were exposed to a phosphoimager screen and the signal was quantified using Image Studio software (LI-COR Biosciences, NE, USA).

### Western blot

HepG2.2.15 or HepG2-NTCP cells were lysed with Laemmli buffer (Bio-Rad, PA, USA). Cell lysates were separated with 12% precast polyacrylamide gels and Tris/Glycine/SDS running buffer (Bio-Rad). Following protein transfer, PVDF membranes were incubated with primary antibody followed by secondary antibody, developed with Clarity™ Western ECL Substrate (Bio-Rad) and imaged by the iBright Imaging Systems (ThermoFisher Scientific). The primary antibodies used in the present study are listed as below, anti-HBc antibody (Dako cat. no. B0586, United Kingdom), anti-PAPD5 antibody (Atlas Antibodies cat. no. HPA042968, Bromma, Sweden), and anti-beta Actin antibody (Abcam cat. no. ab8227, Cambridge, United Kingdom).

### Particle gel for viral nucleocapsid analysis

A particle gel assay was carried out as described previously (44). Secreted viral particles and intracellular viral nucleocapsid from lysed HepG2.2.15 cells were fractionated through nondenaturing 1% agarose gel electrophoresis and blotted to a nitrocellulose filter. To detect HBV core antigens, membranes were probed with antibody recognizing core protein (Dako cat. no. B0586). For the detection of HBV DNA, the membrane was probed with an α-^32^P-UTP labeled HBV minus-strand-specific riboprobe.

### Encapsidated pgRNA

HepG2.2.15 or HepAD38 (45) cell lysates were digested with micrococcal nuclease (20 U/ml) at 37 °C for 30 min to remove unprotected nucleic acids. Core particles were precipitated with 35% PEG-8000 and the associated encapsidated pgRNA was extracted with TRI Reagent™ Solution (Invitrogen™, CA, USA).

### Poly(A) tail-length analysis of HBV transcripts

Poly(A) tails of HBV transcripts were measured with the Poly(A) Tail-Length Assay Kit (Thermofisher) as per manufacturer’s instructions. Detailed methodology and data analysis are provided in the Supporting Information.

### PAPD5 and PAPD7 ATP depletion assay

Reactions were carried out in 10 μl of the reaction mixture (12.5 nM of purified PAPD5 (186-518 a.a.) or PAPD7 (226-558 a.a.) in a buffer containing 10 mM Tris-HCl (pH 8.0), 100 mM KCl, 5mM MgCl2, 250 nM CALM1 (RNA substrate 5′- GCCUUUCAUCUCUAACUGCGAAAAA AAAAA −3′), 750 nM ATP, 0.1 mM EDTA, 1mM TCEP and 0.002% NP-40). Remaining ATP was readout after 3 h incubation using Kinase-Glo® Luminescent Kinase kit following manufacturer’s instructions (Promega, WI, USA).

### In vivo antiviral activity in a mouse model of HBV

Evaluation of *in vivo* antiviral efficacy using AAV-HBV-infected mouse experiments were conducted at Arbutus Biopharma (Burnaby, Canada) within a Canadian Council on Animal Care-accredited Animal Care and Use Program. Male C57BL/6J mice expressed an HBV genotype D variant from an AAV vector. In the assessment of antiviral activity in the adeno-associated viruses (AAV)-HBV-infected mice, statistically significant difference (p<0.05) from vehicle control was determined using one-way ANOVA (Dunn’s multiple comparisons test). Detailed methodology including baseline HBV values are provided in the Supporting Information.

## Authors’ contributions

FL and MG conceived and designed the research. FL, ACHL, FG, ASK, HMS, AM, LB, XW, SC, SGK and AGC performed the research. All authors analyzed the data. FL, ACHL and MG wrote the paper, and AGC, DG, BDD, RR (designed plasmids), AL and MJS revised the paper.

## Acknowledgement

We thank Ingrid Graves, Agnes Jarosz, Chris Pasetka and Alice Li for the analysis of *in vivo* data. Thank Jorge Quintero for scaling up the production of cmpdA. Thank Tianlun Zhou for providing HBsAg-encoded adenoviruses.

## Supporting information

**Fig. S1. Antiviral activity of AB-452 in an AAV-HBV-transduced mouse model.** Animals received AB-452 at 0.1, 0.3, 1 mg/kg or vehicle orally twice daily for 7 days. Effect of AB-452 on the production of (A) serum HBsAg, (B) serum HBV DNA, (C) intrahepatic HBsAg, (D) total HBV RNA and (E) 3.5 kb HBV pgRNA on day 7 post-treatment. (F) Effect of AB-452 on body weight through the 7-day treatment. Data represent group mean (*n* = 5) ± SD. Statistically significant difference (p<0.05) from vehicle control was determined using one-way ANOVA (Dunn’s multiple comparisons test) and is denoted by an asterisk (*).

**Fig. S2. Chemical structure of the HBV capsid inhibitor, cmpdA.**

**Fig. S3. AB-452 reduces HBV RNA levels dose- and time-dependently.** (A) Levels of intracellular pgRNA and sRNA in HepG2.2.15 cells treated with increasing concentrations of AB-452 (0.14 to 1000 nM) for 48 h. (B) Time course analysis of HBV RNAs from cells treated with and without 70 nM AB-452. Total intracellular RNA was extracted from cells harvested at 4, 8, 12, 24 and 48 h time points post-treatment. HBV pgRNA and sRNA were analyzed by Norther blotting with ribosomal RNAs as loading control.

**Fig. S4. HBV RNA poly(A) tail is shortened by AB-452 treatment.** Total RNA was tagged with a poly-G/I tail at the 3’ end and reverse transcribed by poly-G/I specific primer. Both HBV and β-actin mRNA poly(A) tails were specifically amplified using one gene specific primer and the universal primer that anneals to the G/I tail. The obtained amplicon product was resolved on a 2% agarose gel. Gene specific PCR (GSP) was used as loading control. The poly(A) tail length of β-actin mRNA served as the negative control as β-actin mRNA was not affected by AB-452 treatment.

**Fig. S5. SLα deletion reduces sensitivity to AB-452**. H133 and H133_dSLα were transfected into Huh-7 cells and treated with either ETV (1 μM), AB-452 (0.1 μM), or ARB-169451 (0.1 μM) for 5 days. Effect of compounds on HBV was monitored by measuring levels of HBsAg in supernatant and normalized to untreated controls (DMSO). Data represent average values ± standard deviations from at least three independent experiments.

**Fig. S6. Shortening of HBV RNA poly(A) tails by AB-452 treatment or SLα deletion.** Total RNA was tagged with a poly-G/I tail at the 3’ end and reverse transcribed (RT) using a primer specific to the poly-G/I tail. Both HBV and β-actin mRNA poly(A) tails were amplified using a gene specific primer and a universal primer that anneals to the G/I tail. The obtained amplicons were resolved in a 2 % agarose gel. Gene specific PCR (GSP) was used as loading control. The poly(A) tail length of β-actin mRNA served as the negative control as β-actin mRNA was not affected by AB-452 treatment. * labels the potential read-throughs.

**Fig. S7. Confirmation of the *PAPD5*, *PAPD7* and *ZCCHC14* CRISPR-Cas9 mediated knockouts by DNA sequencing.** (A) Insertion-deletion mutations (INDELs) are annotated with the bps of insertion (+) or deletion (−) on alleles, in which “/” is used to separate INDELs among different alleles. The regions targeted by gRNAs are highlighted in black. INDELs are detected by sequencing trace analysis with CAT tool (CRISPR analysis tool). (B) PAPD5 and ZCCHC14 were detected with the indicated antibodies in the Western blots. The * asterisk indicates a cross-reacting band.

**Fig. S8. Knockout of *PAPD5/7* and ZCCHC14 shortened and desensitized HBV RNA poly(A) tail to AB-452.** The poly(A) tail length of HBV sRNA was measured in the *PAPD5/7* single or double KO and *ZCCHC14* KO cell clones treated with and without AB-452 for 5 days. Total RNA was tagged with a poly-G/I tail at the 3’ end and reverse transcribed (RT) using a primer specific to the poly-G/I tail. Both HBV and β-actin mRNA poly(A) tails were amplified using a gene specific primer and a universal primer that anneals to the G/I tail. The obtained amplicons were resolved in a 2 % agarose gel. Gene specific PCR (GSP) was used as loading control. The poly(A) tail length of β-actin mRNA served as the negative control as β-actin mRNA was not affected by AB-452 treatment.

**Fig. S9. PAPD5 is more efficient than PAPD7 to extend poly(A) tail.** Processivity of PAPD5 and PAPD7 poly(A) extension was evaluated in the enzymatic assay in the time course studies. Compared to the PAPD7, PAPD5 was more efficient to extend poly(A) tail on the RNA substrate as demonstrated by measuring the remining ATP in the assay. It is the representative result of two repeated experiments with different time points selected.

